## Supplement for "Sequence-to-sequence translation from mass spectra to peptides with a transformer model"

| | MassIVE-KB count | MassIVE-KB selected | PROSPECT count | PROSPECT selected |
|---|---|---|---|---|
| A | 24910 | 24910 | 540777 | 25090 |
| C | 11473 | 11473 | 90772 | 38527 |
| D | 44654 | 44654 | 267716 | 5346 |
| E | 42977 | 42977 | 603469 | 7023 |
| F | 38183 | 38183 | 884261 | 11817 |
| G | 21640 | 21640 | 344670 | 28360 |
| H | 166398 | 50000 | 357268 | 0 |
| I | 12984 | 12984 | 437698 | 37016 |
| K | 16289160 | 50000 | 960538 | 0 |
| L | 50002 | 50000 | 2264210 | 0 |
| M | 19405 | 19405 | 388750 | 30595 |
| N | 40867 | 40867 | 294550 | 9133 |
| P | 9685 | 9685 | 114504 | 40315 |
| Q | 33572 | 33572 | 542063 | 16428 |
| R | 13589301 | 50000 | 1135502 | 0 |
| S | 28389 | 28389 | 457519 | 21611 |
| T | 15845 | 15845 | 443535 | 34155 |
| V | 26011 | 26011 | 774624 | 23989 |
| W | 4261 | 4261 | 341456 | 45739 |
| Y | 35180 | 35180 | 1368308 | 14820 |
| Total | 30504897 | 610036 | 12612190 | 389964 |

Table S1: **Creating a non-enzymatic dataset by sampling from PROSPECT and MassIVE-KB.** PROSPECT was first downsampled to include at most 100 PSMs per peptide sequence. MassIVE-KB and PROSPECT were then segregated by C-terminal amino acid, and we randomly selected from each category from MassIVE-KB, supplementing as necessary from PROSPECT to obtain 50,000 PSMs per terminal amino acid.

| PRIDE | Species | Uniprot | Files | Spectra | PSMs | Peptides | precursor | fragment |
|---|---|---|---|---|---|---|---|---|
| PXD005025 | *Vigna mungo* | UP000087766 | 24 | 932848 | 108514 | 12001 | 20 | 0.05 |
| PXD004948 | *Mus musculus* | UP000000589 | 13 | 306786 | 25541 | 5899 | 10 | 0.05 |
| PXD004325 | *Methanosarcina mazei* | UP000033058 | 72 | 3728183 | 267333 | 15925 | 10 | 0.05 |
| PXD004565 | *Bacillus subtilis* | UP000001570 | 106 | 4336428 | 1358337 | 30786 | 30 | 0.05 |
| PXD004536 | *Candidatus endoloripes* | UP000094849 | 11 | 2272023 | 82290 | 8392 | 20 | 0.05 |
| PXD004947 | *Solanum lycopersicum* | UP000004994 | 60 | 603506 | 178413 | 49745 | 15 | 0.05 |
| PXD003868 | *Saccharomyces-cerevisiae* | UP000002311 | 27 | 1477397 | 585593 | 19720 | 20 | 0.05 |
| PXD004467 | *Apis mellifera* | UP000005203 | 17 | 823169 | 194281 | 21559 | 20 | 0.05 |
| PXD004424 | *H. sapiens* | UP000005640 | 26 | 684821 | 44604 | 11289 | 20 | 0.02 |
| Total | | | 343 | 15,165,161 | 2,844,906 | 175,316 | | |

Table S2: **The nine-species benchmark.** The final two columns specify the precursor window size (in ppm) and fragment bin size (in Da) used in the database search step. No reference proteome is available for *Vigna mungo*, so the proteome for the closely related species *Vigna radiata* was used instead.

| In human proteome | In Casanovo peptide | BLOSUM score | Count |
|---|---|---|---|
| L | V | 1 | 749 |
| V | L | 1 | 650 |
| E | Q | 2 | 467 |
| N | D | 1 | 437 |
| R | K | 2 | 371 |
| E | H | 0 | 343 |
| E | D | 2 | 338 |
| L | K | -2 | 314 |
| L | M | 2 | 304 |
| L | F | 0 | 301 |

Table S3: **The most common amino acid swaps have positive BLOSUM scores.** We found that the top ten most common single amino acid substitutions that can be explanied with a single nucleotide polymorphism detected by Casanovo are enriched for positive BLOSUM scores.
